# Targeting mitochondria mitigates chemotherapy-induced bone marrow dysfunction

**DOI:** 10.1101/2025.07.18.665515

**Authors:** Tim V. D. van Tienhoven, Johanna Lepland, Anne de Snaijer, Akin Bucakci, Wout Maaskant, Gregory van Beek, Eric M J Bindels, Marc H. G. P. Raaijmakers, Tariq Enver, Valgardur Sigurdsson, Els Mansell

## Abstract

Chemotherapy has revolutionized cancer treatment but its long-term impact on healthy tissues, particularly the rapidly dividing hematopoietic system, remains a significant concern. We show that the chemotherapeutic agent 5-fluorouracil (5-FU) causes significant long-term defects of the hematopoietic system that mimics ageing, including myeloid lineage skewing and hematopoietic stem cell (HSC) dysfunction. Importantly, chemotherapy exposed HSCs remain in an inflammatory state coupled with mitochondrial dysfunction, both of which are implicated in ageing and MDS development. Remarkably, five days of MitoQ treatment fully reversed 5-FU-induced myeloid skewing and caused significant recovery of HSC transcriptome and function in a stable manner. Thus, our results demonstrate that repeated chemotherapy induces long-term bone marrow dysfunction driven by metabolically unfit HSCs that can be pharmacologically rescued. This work opens up novel avenues to explore supportive treatment for patients undergoing any type of myelo-ablative chemotherapy.

**Highlights:**

- Commonly used chemotherapy drugs cause stable lineage-specific cytopenias despite recovery of total blood counts.
- 5-FU-induced hematopoietic dysfunction phenocopies premature hematopoietic ageing.
- Chemotherapy-induced changes stem from HSCs that are self-renewal competent but differentiation incompetent.
- Mitochondrial-targeted treatment rescues 5-FU-induced myeloid bias through correction of HSC transcriptome and function.

## Introduction

Hematopoietic stem cells (HSCs) occupy the apex of the hematopoietic differentiation hierarchy and replenish all blood lineages throughout life (1, 2). Hematopoietic homeostasis is achieved by appropriately balancing HSC self-renewal and differentiation (3, 4). HSCs are predominantly quiescent under homeostatic conditions, yet they retain the capacity to rapidly respond to hematological stressors such as infections or exposure to cytotoxic treatments (4–9). Cytotoxic drugs, including the chemotherapeutic agent fluorouracil (5-FU), induce myelosuppression by broadly ablating rapidly dividing hematopoietic cells (10). Quiescent HSCs are spared from 5-FU-induced cell killing and are subsequently activated to regenerate hematopoiesis (11, 12).

HSC activation and return to dormancy requires tight metabolic control, to meet the energetic demands required for regeneration whilst preserving HSC integrity (13–17). Mitochondria are emerging as critical organelles driving HSC behaviour, and mitochondrial dysfunction is linked to ageing and transformation (17–23).

In clinical practice, hematopoietic recovery between chemotherapy cycles is critical for treatment continuation and is typically assessed by monitoring only neutrophil and platelet counts (24). Although partially indicative of recovery, these parameters do not capture incomplete recovery or long-term effects on other mature lineages, progenitors, or HSCs. Incomplete hematopoietic recovery does not only impact on the patient’s quality of life and treatment efficacy, but is also an independent risk factor for non-relapse mortality (25, 26).

Here, we explore the long-term effects of clinically relevant chemotherapeutic regimens on the hematopoietic system, with a specific focus on HSC recovery in terms of phenotype, metabolism, transcriptional state and function. We show that despite recovery of neutrophil and platelet counts, chemotherapy induces long-term hematopoietic damage, as indicated by persisting myeloid-biased peripheral blood output and loss of HSC function. Chemotherapy-induced hematopoietic dysfunction mimics natural ageing and mitochondrial deflation is known to be causative of that. To rescue this chemotherapy-induced phenotype we tested whether mitochondrial potentiation treatment, previously shown by us to functionally restore HSCs from aged mice, could be beneficial in this context (20). Strikingly, we show that strategic treatment with the mitochondrial-targeted drug MitoQ improves HSC function and reverses myeloid-biased peripheral blood output after 5-FU. This work opens up novel avenues to make cancer treatment more effective and better tolerated through mitochondrial supportive therapy.

## Results

### Chemotherapy induces premature hematopoietic ageing

To explore whether multi-cycle chemotherapy treatment induces long-term alterations to the hematopoietic system, wild type mice received three intravenous administrations of 150mg/kg of 5-FU at three-week intervals (**Figure 1A**). This interval is routinely used in clinical settings and coincides with complete numerical recovery of white blood cell counts, neutrophil, platelet counts (**Figure S1A/B**), and near complete recovery of BM cytokine levels (**Figure S1C**). However, we observe that despite this apparent recovery, peripheral blood (PB) output remains myeloid-skewed. 5-FU-treated mice show hampered recovery of red blood cells and B cells and exaggerated production of neutrophils and platelets (**Figure 1B/C**). Mirroring the PB observations, the bone marrow (BM) of 5-FU exposed mice contains a significantly smaller proportion of common lymphoid progenitors (CLP) compared to control mice, suggesting that the shift towards the myeloid lineage is initiated high up in the hematopoietic hierarchy (**Figure 1D**). To address whether this myeloid-biased output is attributable to alterations within the hematopoietic stem cell (HSC) compartment, we performed detailed analysis of immunophenotypically defined HSCs (Lin^-^ c-Kit^+^ Sca-1^+^ CD150^+^ CD48^-^) after chemotherapy (**Figure 1E**) (27). We observe a significant increase in the frequency of immunophenotypic HSCs, that show elevated CD150 expression. Interestingly, CD150 expression distinguishes between myeloid-biased HSCs (My; CD150^high^) and lineage-balanced HSCs (Ly; CD150^low^) (28), suggesting that the expanded HSC pool is skewed towards a myeloid-biased lineage output, a phenomenon commonly associated with ageing (29). To further scrutinize changes to HSC heterogeneity, we used the marker CD34 to discriminate between long-term (CD34^-^) and short-term (CD34^+^) HSCs (30). Mice that have been exposed to 5-FU treatment, have a significant increase in long-term HSCs. We could furthermore see identical changes in the BM and HSC pool after just one treatment with the anthracycline drug doxorubicin (doxo) (**Figure S1D-H**), suggesting that these changes are inherent to recovery from cytotoxicity-induced myelosuppression, rather than arising from drug-specific pathways. To evaluate whether the immunophenotypically altered HSCs exhibit impaired functional capacity, we performed transplantation experiments (**Figure 1F**). Four hundred HSCs harvested from chemotherapy-treated or untreated mice were transplanted with 200.000 SJL whole bone marrow support cells into lethally irradiated SJL mice. HSCs from single Doxo-injected mice showed markedly reduced BM reconstitution potential (**Figure S1F-H**). HSCs from 5-FU exposed animals exhibited extremely low chimerism in the peripheral blood (**Figure 1G**) and BM (**Figure 1H**) compared to HSCs from untreated controls, despite showing comparable and near complete HSC pool reconstitution (**Figure 1H**). These data indicate that multi-cycle 5-FU treatment results in HSCs that are self-renewal competent but differentiation incompetent. We are intrigued by the possibility that these changes persist and cause long-term hematopoietic damage.

**Figure 1.**
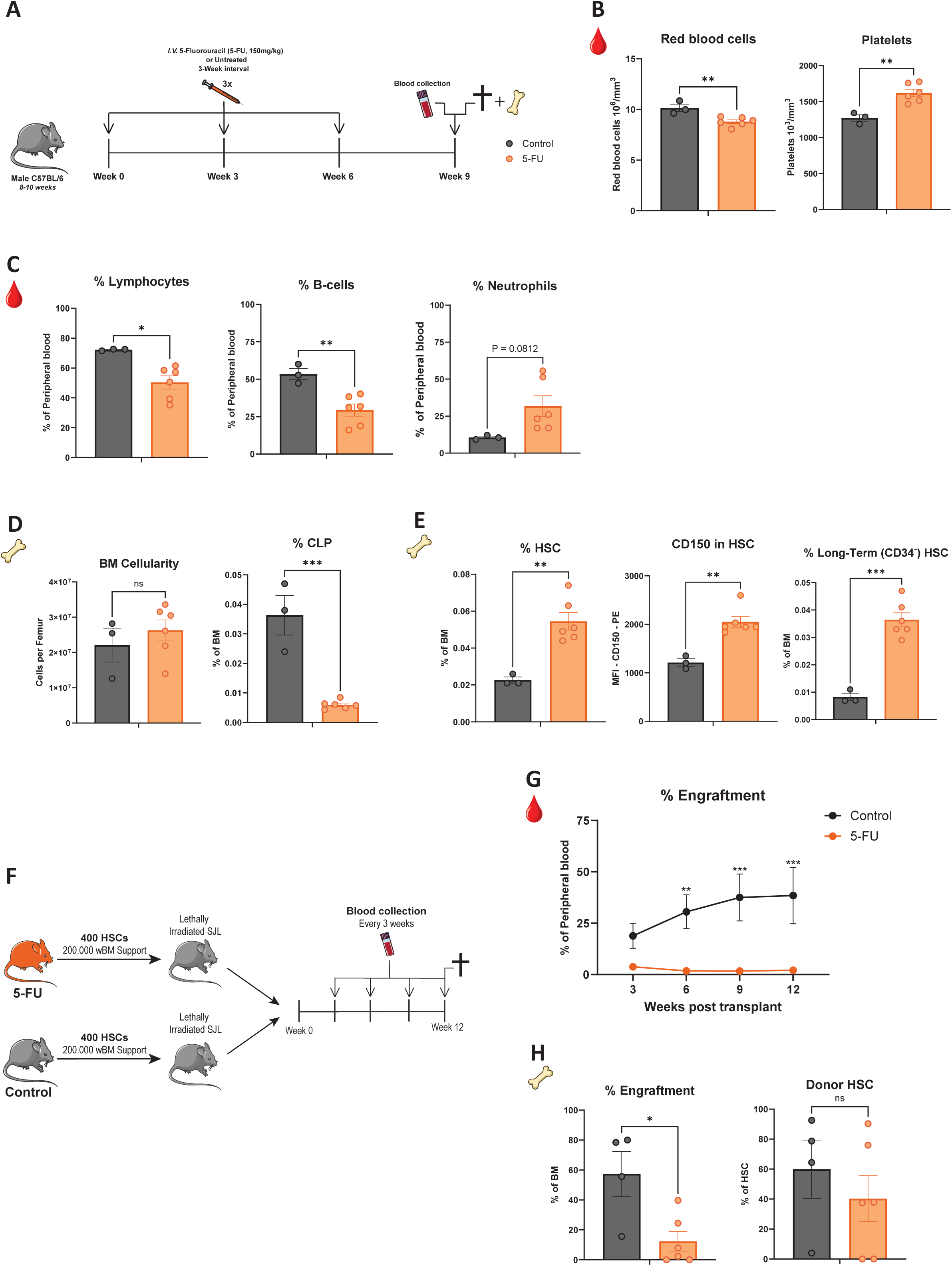
Chemotherapy induces “clinically hidden” premature hematopoietic ageing. (A) Experimental overview. (B) Scil vet ABC plus counter analysis of red blood cells (Left) and platelets (Right) in peripheral blood (PB) of control (N=3, black) or 5-FU exposed (N=6, orange) animals 3 weeks post-chemotherapy treatment. (C) Left: Scil vet ABC plus counter analysis of percentage lymphocytes in PB. Percentage B cells (B220^+^) (Middle) and neutrophils (Gr-1^+^ CD11b^+^ CD115^-^) (Right) in PB 3 weeks post-chemotherapy treatment. (D) Left: BM cellularity counts and percentage CLP (Lineage^-^ c-Kit^intermediate^ Sca-1^intermediate^ CD127^+^ FLT3) of total BM (Right) (E) Left: Percentage of HSCs (Lin^-^ C-kit^+^ Sca-1^+^ CD150^+^ CD48^-^) of total BM. Middle: Quantification of median fluorescence intensity of CD150 in HSCs. Right: Percentage of Long-Term HSCs (Lin^-^ C-kit^+^ Sca-1^+^ CD150^+^ CD48^-^ CD34^-^) of total BM. (F) Transplantation setup. (G) Total PB engraftment in recipients receiving 5-FU exposed (N=6, orange) or control (N=6, black) HSCs. (H) Total BM engraftment (Left) and contribution of donor HSC to total HSC pool (Right) 12 weeks after transplantation. Results are represented as mean ± SEM. ^∗^p < 0.05; ^∗∗^p < 0.01; ^∗∗∗^p < 0.001. Statistical analysis used: unpaired two-tailed Student’s *t*-test for two group comparisons and two-way ANOVA followed by Šídák’s test for multiple comparisons.

### Chemotherapy induces long-term hematopoietic damage

To evaluate the long-term persistence and potential exacerbations of these alterations following chemotherapy, mice were subjected to the same multi-cycle treatment regimen and analyzed three months, instead of three weeks, post-treatment (**Figure 2A**). PB analysis at 3, 6, and 12 weeks post-chemotherapy revealed a complete numerical recovery of white blood cells, red blood cells, platelets (**Figure S2A**), and BM counts (**Figure S2B**). Despite this ‘complete hematopoietic recovery’, we observe a sustained myeloid-biased PB output. Notably, by 12-weeks post-treatment, 5-FU exposed mice exhibited significantly fewer circulating B cells and increased levels of neutrophils compared to control mice (**Figure 2B**). In addition, there was no recovery of BM composition with persistent loss of CLPs (**Figure 2C**) and a significant expansion of the HSC pool (**Figure 2D**). Critically, this pool of expanded long-term HSCs displayed major mitochondrial alterations, as both total mitochondrial content (Mitotracker green [MTG]) and mitochondrial membrane potential (Tetramethylrhodamine methyl ester [TMRM]), exhibit a total reduction of 50% (**Figure 2E**). This indicates that these long-term HSCs are metabolically deflated, a state previously associated with hematopoietic ageing and stem cell dysfunction (20).

**Figure 2.**
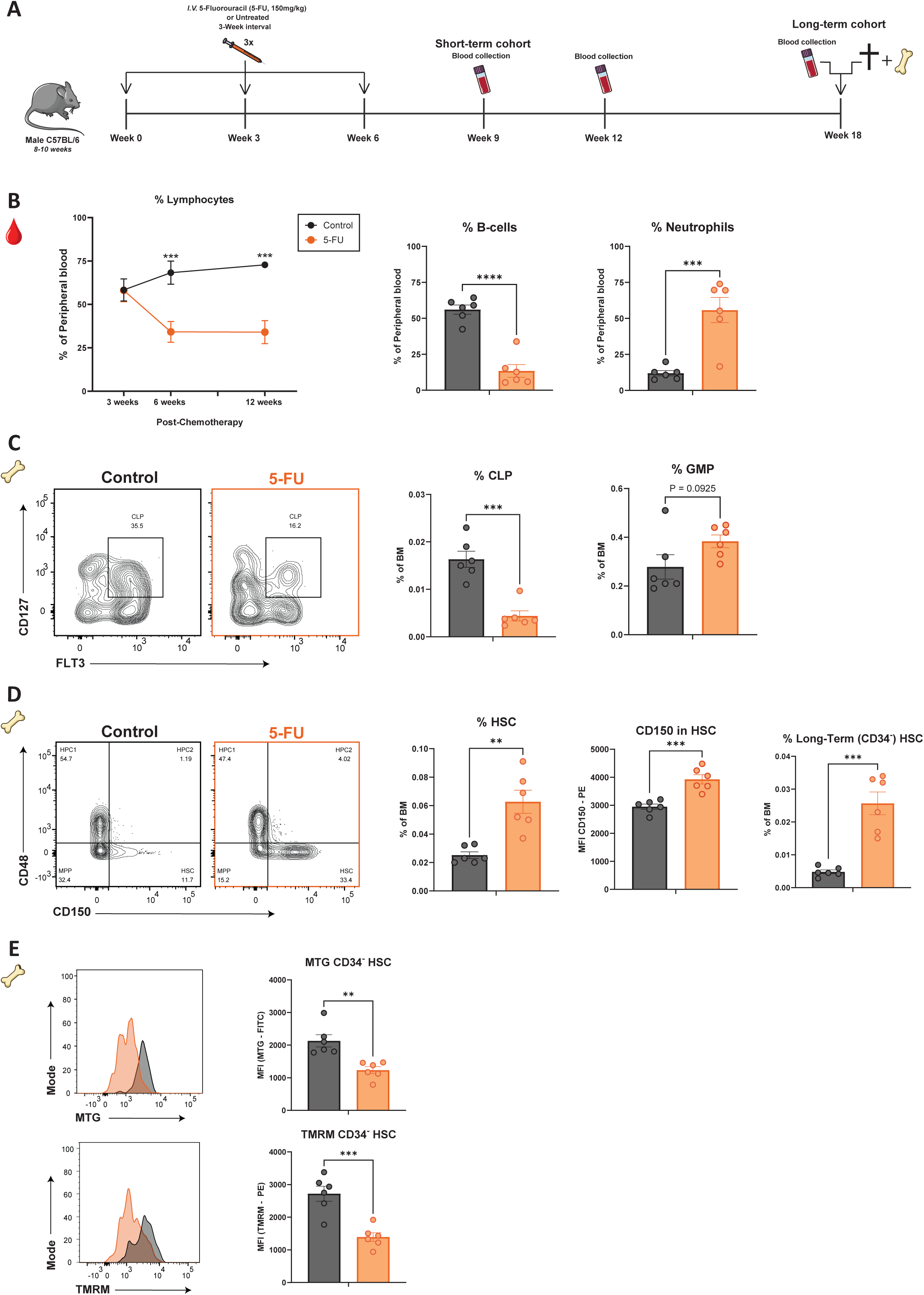
Chemotherapy induces long-term hematopoietic damage. (A) Experimental overview. (B) Left: Scil vet ABC plus counter analysis of percentage lymphocytes in peripheral blood (PB) of control (black, N=6) and 5-FU exposed animals (orange, N=6). Percentage B cells (B220^+^) (Middle) and neutrophils (Gr-1^+^ CD11b^+^ CD115^-^) (Right) in PB 12 weeks post-chemotherapy treatment. (C) Left: Representative FACS plots of common-lymphoid progenitor (CLP, Lineage^-^ c-Kit^intermediate^ Sca-1^intermediate^ CD127^+^ FLT3^+^). Right: percentage CLP and granulocyte-monocyte progenitor (GMP, Lineage^-^ C-Kit^+^ Sca-1^-^ CD16/32^+^ CD34^+^) of total BM 12 weeks post-chemotherapy. (D) Representative FACS plots of hematopoietic stem cells (Lin^-^ C-kit^+^ Sca-1^+^ CD150^+^ CD48^-^). Middle: Frequency of phenotypic HSC in total BM and analysis of median fluorescence intensity of CD150 in HSC. Right: percentage Long-Term HSCs (Lin^-^ C-kit^+^ Sca-1^+^ CD150^+^ CD48^-^ CD34^-^) of total BM 12 weeks post-chemotherapy. (E) Representative FACS plots and quantifications of mitochondrial mass (MTG; upper) and potential (TMRM; lower) between control and 5-FU exposed animals. Results are represented as mean ± SEM. ^∗∗^p < 0.01; ^∗∗∗^p < 0.001; ^∗∗∗∗^p < 0.0001. Statistical analysis used: unpaired two-tailed student’s *t*-test for two group comparisons and two-way ANOVA followed by Šídák’s test for multiple comparisons.

Collectively, these results show that three administrations of 5-FU treatment induce a persistent myeloid-bias in PB, accompanied by phenotypic alterations in HSCs that resemble a prematurely aged state.

### HSCs are persistently transcriptionally altered by 5-FU treatment

We next sought to understand whether the persistent phenotypic and metabolic changes were reflected in a chronic transcriptional state change of HSCs. To this end, we performed RNA sequencing of HSCs after repeated 5-FU exposure. HSCs were isolated from mice after an extended recovery period of 12 weeks to make sure that the phenotype was not transient (**Figure 3A**). Principal-component analysis (PCA) reveals a clear separation of control and 5-FU exposed HSCs (**Figure 3B**), indicating long-term transcriptional reprogramming of HSCs. We next explored the underlying basis of this transcriptional difference, by focusing on the significantly differentially expressed genes (DEGs, log2 fold change ≥ 0.5, adjusted *P*-value ≤ 0.05). Notably, 760 genes are significantly altered by 5-FU exposure, of which 381 are upregulated and 379 downregulated (**Figure 3C**). Our data indicate persistent downregulation of critical regulators of HSC function (*Gprc5c, Igf2bp2, CD86)* and persistent upregulation of genes linked to a decline in HSC function or ageing (*CD244a*, *CD48*, *Ly6a*, *Neo1*, *Pim1, Selp, tgfbi)* after 5-FU exposure (**Figure 3D**) (28–37). This suggests that fundamental stem cell programs remain dysregulated long-term after 5-FU exposure, despite the attainment of a numerically defined steady-state hematopoiesis. We next performed gene set enrichment analysis (GSEA) to further explore the nature of transcriptional change in an unbiased manner. In line with our observation of metabolic deflation **(Figure 3E / S3A)**, we found significant downregulation of the Mitocarta gene set, the most extensive inventory of genes involved in mammalian mitochondrial proteins and pathways, in 5-FU-exposed HSCs (**Figure S3B)**.

**Figure 3.**
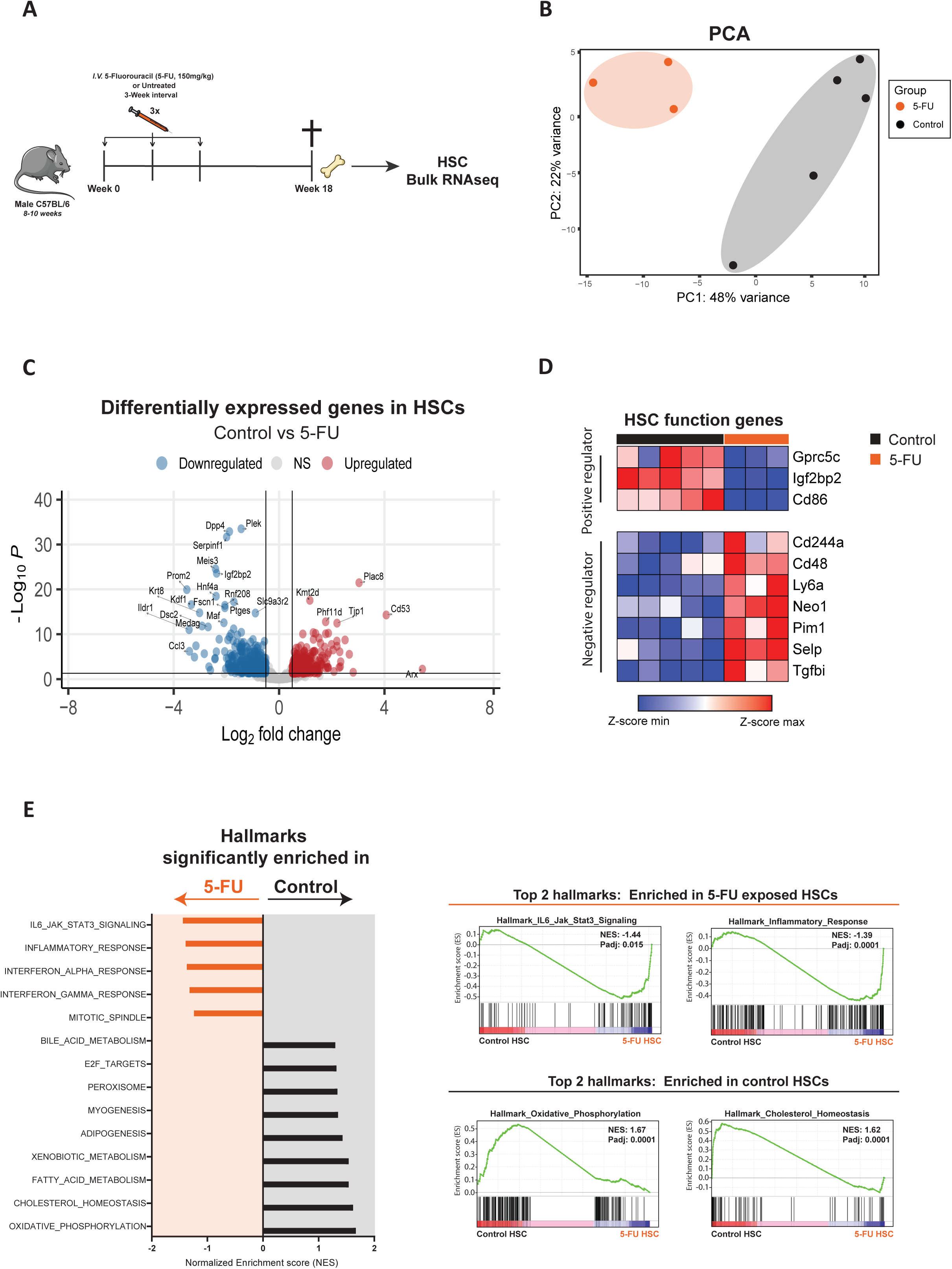
HSCs are persistently transcriptionally altered after 5-FU treatment. (A) Experimental overview. (B) Principal-component analysis (PCA) of HSCs sorted from 5-FU exposed (N=3, orange) and control (n=5, black) animals 12 weeks post-chemotherapy. (C) Volcano plot of differentially expressed genes (DEGs, Log2fold change ≥ 0.5, adjusted *P*-value ≤ 0.05) comparing 5-FU exposed and control HSCs. Upregulated DEGs are depicted in red and downregulated DEGS in blue. (D) Heatmap of selected DEGs involved in HSC function. (E) Gene set enrichment analysis (GSEA) showing top significantly up- or downregulated pathways comparing HSCs of 5-FU exposed and control animals.

Indeed, the most significantly downregulated gene sets after 5-FU encompass metabolic processes including oxidative phosphorylation, cholesterol homeostasis, fatty acid oxidation (FAO), bile acid, acetyl-CoA and pyruvate metabolism (**Figure 3E and S3C**). These metabolic pathways play a pivotal role in HSC homeostasis, and their disturbance may influence HSC survival and fate decisions (38). Importantly, 5-FU exposed HSCs were significantly enriched for inflammatory signatures, including the hallmarks for inflammatory response, IFN-γ response, and IFN-α response. This intrinsic retention of an inflammatory state long after cytotoxic exposure is reminiscent of sterile ‘inflammageing’ and may have implications for stem cell fitness, DNA damage and clonal advantage of mutated HSCs (39–46). These data show that repeated 5-FU exposure induces a retained transcriptional state of sterile inflammation and metabolic impairment.

### MitoQ treatment re-establishes lineage balanced blood production after 5-FU

Our previous work has shown that mitochondrial-targeted treatment can alleviate multiple aspects of hematopoietic ageing. Since the 5-FU induced changes mimic premature hematopoietic ageing, we tested whether a similar treatment strategy was applicable in the context of chemotherapy-induced bone marrow dysfunction.

To this end, 5-FU exposed mice were treated with the drug mitoquinol mesylate (MitoQ), a mitochondrial-targeted antioxidant composed of a triphenylphosphonium cation (TPP^+^) linked to a ubiquinone moiety via a 10-carbon chain (47, 48). We have shown that MitoQ treatment can fully revert the myeloid-biased blood output in aged mice, by changing quantitative and qualitative transcriptional programs in HSCs (20). We treated mice with 5 consecutive daily injections of MitoQ, 21 days after the final 5-FU treatment (**Figure 4A**). Our aim was to test if MitoQ could rescue the persistent chemotherapy-induced hematopoietic dysfunction. Remarkably, MitoQ treatment caused a significant and complete rescue of total percentage of lymphocytes in the PB of 5-FU exposed mice (**Figure 4B**). The skewed distribution towards neutrophils was completely and stably corrected with one course of MitoQ treatment. MitoQ also accelerated the normalization of exaggerated platelet counts (**Figure S4A**). In further support of MitoQ’s ability to rescue the 5-FU-induced phenotype, we performed H&E staining on the humeri from all animals at the experimental endpoint. We observed a significant increase in adipocyte numbers in the humeri of 5-FU only treated animals (**Figure 4C**), consistent with natural ageing (49, 50). Strikingly, MitoQ treatment at week 9 resulted in fully normalized numbers of fat cells at the 18-week experimental endpoint. This indicates that the hematopoietic restoration induced by MitoQ extends to the bone marrow architecture.

**Figure 4.**
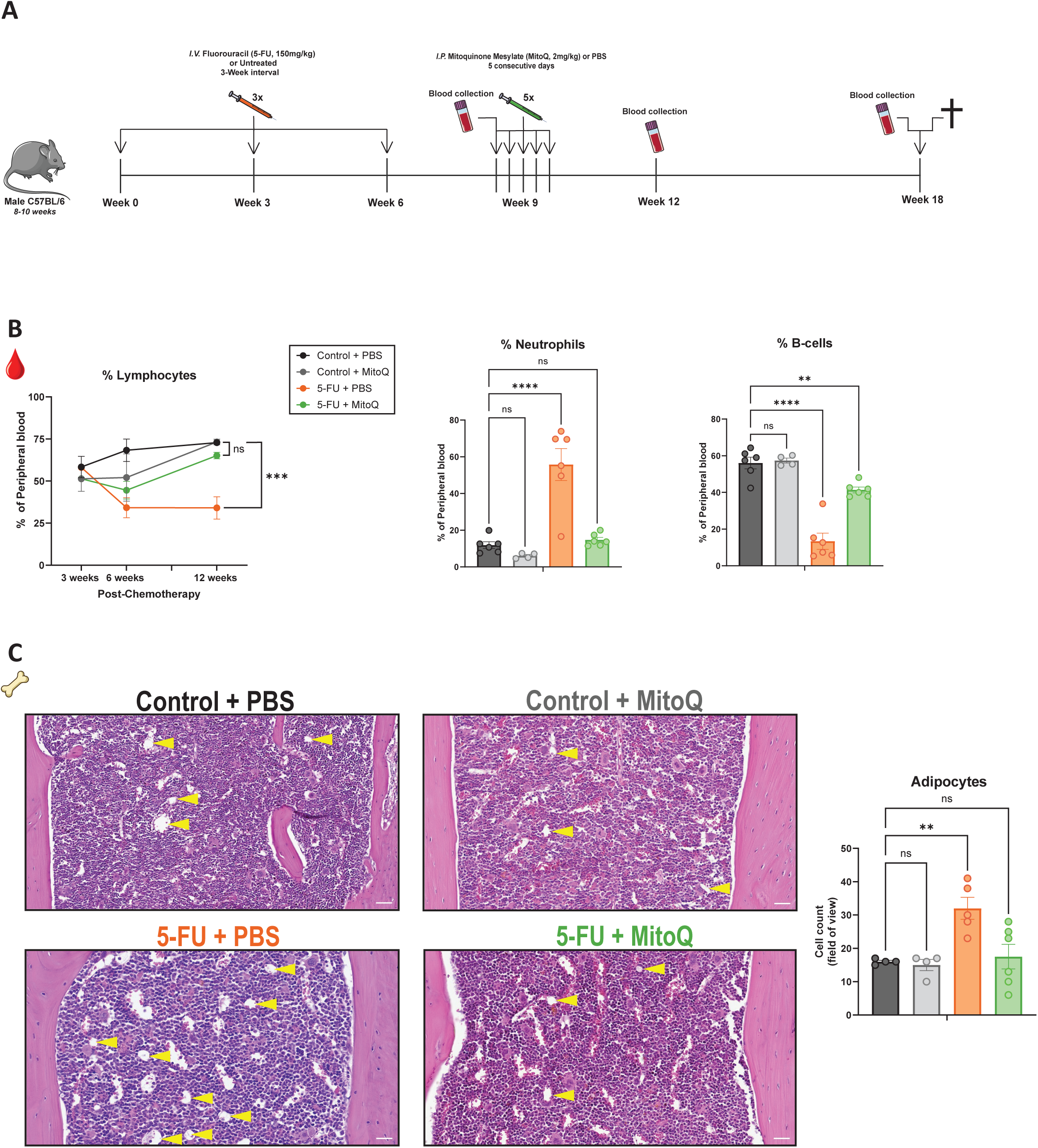
MitoQ treatment re-establishes lineage balanced blood production after 5-FU. (A) Experimental overview. (B) Scil vet ABC plus counter analysis of percentage lymphocytes in peripheral blood (PB) 3, 6, and 12 weeks post-chemotherapy of control + PBS (N=6, black), control + MitoQ (n=4, grey), 5-FU + PBS (N=6, orange) and 5-FU + MitoQ (n=6, green). Percentage B cells (B220^+^) (Middle) and neutrophils (Gr-1^+^ CD11b^+^ CD115^-^) (Right) in PB 12 weeks post-chemotherapy treatment. (C) Left: Representative H&E images (White scale bar indicates 0.05 mm) of mouse humeri 12 weeks post-chemotherapy. Yellow arrows indicate adipocytes. Right: quantification of adipocytes. Results are represented as mean ± SEM. ^∗∗^p < 0.01; ^∗∗∗^p < 0.001; ^∗∗∗∗^p < 0.0001. Statistical analysis used: one-way analysis of variance (ANOVA) followed by Dennett’s multiple comparisons test for comparisons to control and two-way ANOVA followed by Šídák’s test for multiple comparisons.

### MitoQ treatment rescues HSC transcriptional programs essential for functional output

We were intrigued by the possibility that MitoQ’s effect manifests in HSCs, explaining the durable nature of the corrected PB output. Therefore, we assessed whether MitoQ treatment also durably corrects the transcriptional state of HSCs exposed to 5-FU. To this end, we performed RNA sequencing of HSCs harvested at the 18-week experimental endpoint. Interestingly, PCA analysis reveals that MitoQ treatment markedly shifts the transcriptional state of 5-FU exposed HSCs towards steady state HSCs isolated from vehicle-treated control animals (**Figure 5A**). To further explore the nature of transcriptional changes, we determined DEGs by comparing HSCs isolated from 5-FU exposed animals with those isolated from 5-FU + MitoQ-treated animals. MitoQ treatment significantly altered 82 genes, of which 36 were upregulated and 46 were downregulated (**Figure S5A/B**). Interestingly, 53 out of the 82 genes, are genes that were significantly altered by 5-FU treatment and now restored by MitoQ treatment (**Figure 5B**). Several of these 53 genes are key functional regulators of HSCs. MitoQ completely rescues expression of HSC dormancy regulator *GPCR5C* and downregulates genes involved in stem cell inflammageing, myeloid bias or dysfunction (*CD244a, Sult1a1, Selp, Cebpd, Tgfbi, Apobec3) (***Figure 5C**) (51–53). Moreover, MitoQ restores the expression of genes involved in metabolic pathways *(Slc16a9, Coq8a, Atp9a).* Beyond DEG analysis, we performed GSEA to further explore the nature of transcriptional changes (**Figure 5D**). GSEA revealed that MitoQ treatment normalized the 5-FU-induced transcriptional enrichment for MYC targets and unfolded protein response (UPR) in HSCs. UPR upregulation is associated with ER- and cellular stress and suggests that protein homeostasis remains deregulated (54–57). In addition, we observed a near significant enrichment (Normalized P-value 0.065) of inflammatory response in the 5-FU exposed HSCs compared to 5-FU + MitoQ-exposed HSCs, indicating a dampening of inflammatory transcriptional signatures by MitoQ treatment. Importantly, MitoQ fully restores expression of the most significantly downregulated pathway after 5-FU: cholesterol homeostasis. It was recently shown that mitochondrial NADPH fuels cholesterol biosynthesis in HSCs in a FAO-dependent manner, supporting HSC function and homeostasis (58).

**Figure 5.**
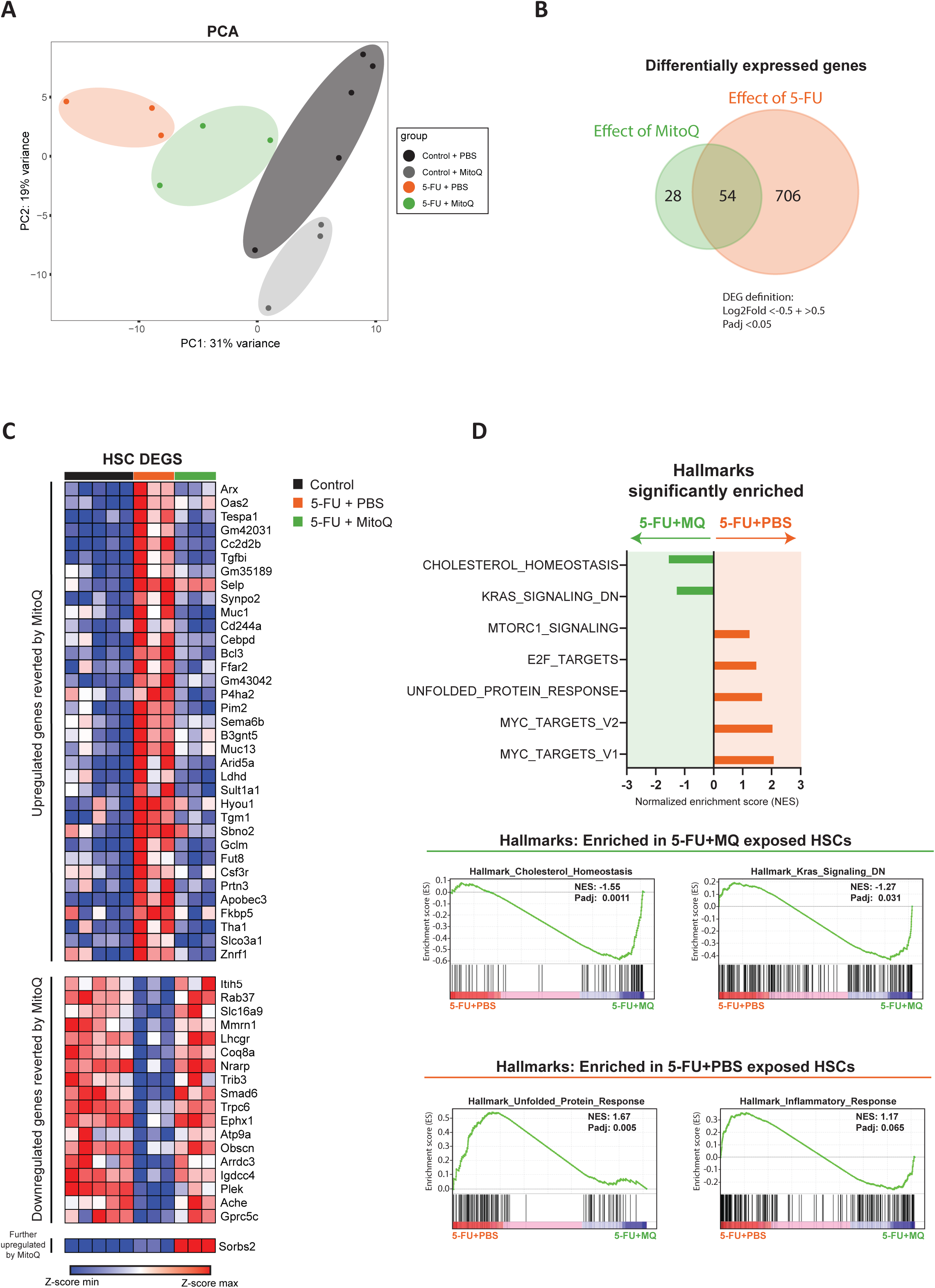
MitoQ treatment rescues transcriptional programs highly relevant for HSC function. (A) Principal-component analysis (PCA) of HSCs sorted from control + PBS (N=5, black), control + MitoQ (n=3, grey), 5-FU + PBS (N=3, orange) and 5-FU + MitoQ (N=3, green) 12 weeks post-chemotherapy. (B) Venn diagram showing the overlap of differentially expressed genes (DEGs, Log2fold change ≥ 0.5, adjusted *P*-value ≤ 0.05) by comparing control + PBS vs 5-FU + PBS (orange) and 5-FU + PBS and 5-FU + MitoQ (green). (C) Heatmap of 54 selected DEGs, of which 53 exhibited reversed expression and 1 showed exacerbated expression following MitoQ treatment. (D) Gene set enrichment analysis (GSEA) showing top significantly up- or downregulated pathways comparing HSCs of 5-FU + PBS and 5-FU + MitoQ exposed animals.

Thus, one treatment regimen with MitoQ 21-days post-chemotherapy elicited HSC intrinsic changes that were maintained approximately 3 months later, with rescue of transcriptional programs highly relevant for HSC function. To test whether this transcriptional rejuvenation would translate to improved HSC function, we performed transplantation experiments.

### MitoQ treatment improves HSCs function and lineage output

To assess the extent to which 5-FU treatment persistently alters HSC function and whether MitoQ could rescue this, we performed transplantation assays. Four hundred HSCs that were isolated three months after exposure to 5-FU, with or without MitoQ treatment, were transplanted in combination with 200.000 BM support cells, into lethally irradiated SJL mice (**Figure 6A**). We found that HSCs from 5-FU exposed mice still exhibited significantly lower engraftment in PB (**Figure 6B**) and BM (**Figure 6C**) up to 15-weeks post-transplantation, compared to HSCs from untreated controls, despite comparable and near-complete donor contribution to the HSC pool (**Figure 6D**). This indicates an accumulation and expansion of self-renewing HSCs in the BM that remain impaired in their differentiation capacity and contribution to blood. In addition, HSCs from 5-FU exposed mice produce significantly less lymphoid output in both PB and BM (**Figure 6B/C**). Notably, B cell output was significantly impaired compared to untreated controls, indicating that 5-FU treatment persistently and intrinsically changes HSC behaviour. MitoQ treatment markedly attenuates these changes, by significantly increasing both total chimerism and B cell output in the PB and BM (**Figure 6B/C)**. Notably, MitoQ normalized the contribution of 5-FU exposed HSCs to the megakaryocyte-erythrocyte progenitor (MEP) population (**Figure 6D**). Thus, MitoQ facilitates partial rescue of both the quantitative and qualitative regeneration potential of HSCs after cytotoxic exposure. Of note, MitoQ treatment alone, in absence of cytotoxic drugs, resulted in modestly increased chimerism upon transplantation without detectable changes to PB or BM composition.

**Figure 6.**
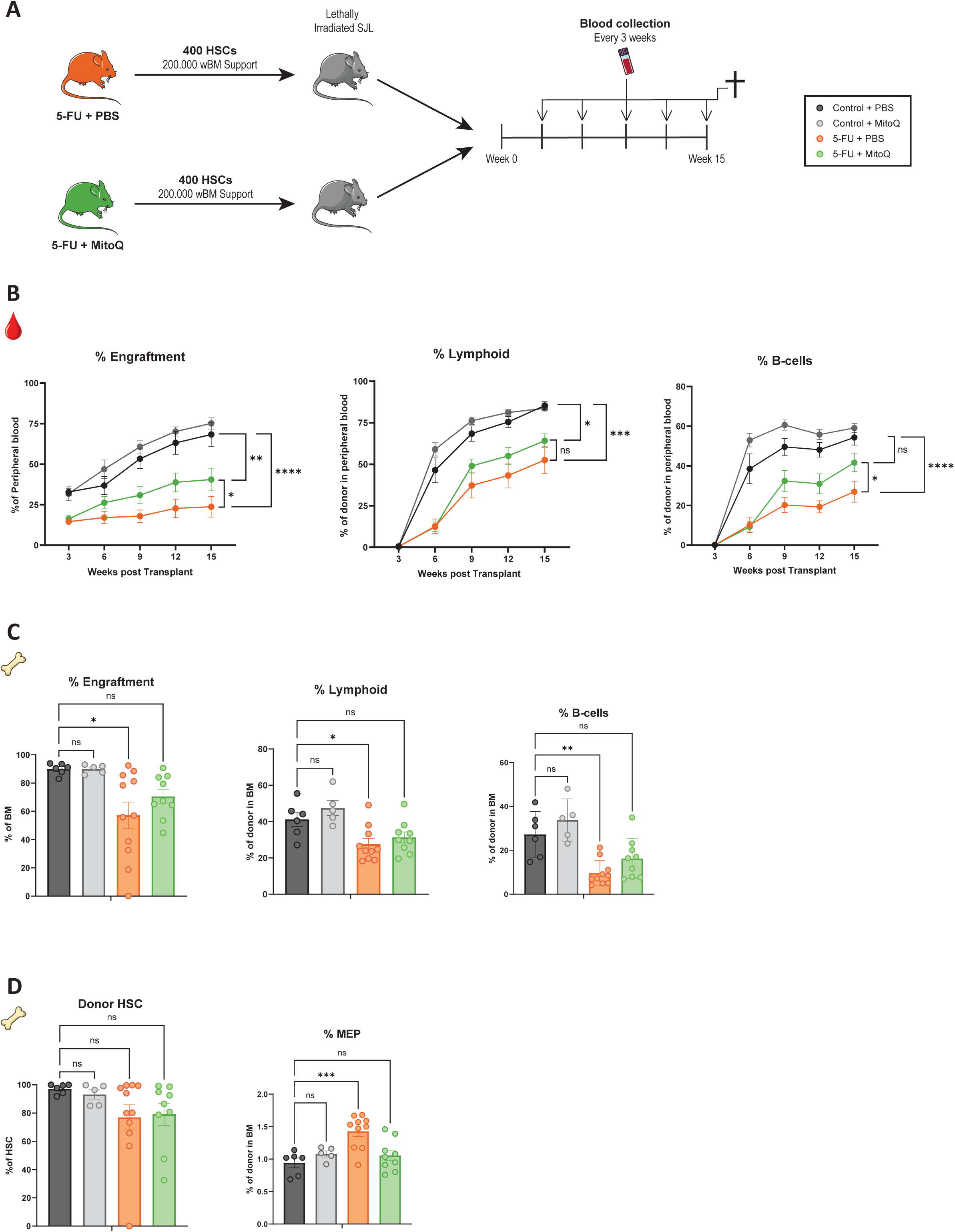
MitoQ treatment improves HSCs function and lineage output. (A) Transplantation setup. (B) Total peripheral blood (PB) engraftment in recipients receiving HSCs of control + PBS (N=6, black), control + MitoQ (n=5, grey), 5-FU + PBS (N=11, orange) and 5-FU + MitoQ (N=9, green) (Left). Percentage lymphoid (Gr-1^-^ CD11b^-^) (Middle) and B cells (B220^+^) (Right) in PB (C) Total BM engraftment (Left) control + PBS (N=6, black), control + MitoQ (n=5, grey), 5-FU + PBS (N=11, orange) and 5-FU + MitoQ (N=9, green). Middle:percentage lymphoid (Gr-1^-^ CD11b^-^), B cells of donor in BM (B220^+^) (Right) control + PBS (N=6, black), control + MitoQ (n=5, grey), 5-FU + PBS (N=10, orange) and 5-FU + MitoQ (N=9, green). (D) (D) Contribution of donor HSC to total HSC pool (Right). Percentage MEP (Lin^-^ C-Kit^+^ Sca-1^-^ CD34^-^ CD16/32^-^) of donor in BM. Results are represented as mean ± SEM. ^∗^p < 0.05; ^∗∗^p < 0.01; ^∗∗∗^p < 0.001; ^∗∗∗∗^p < 0.0001. Statistical analysis used: one-way analysis of variance (ANOVA) followed by Dennett’s multiple comparisons test for comparisons to control and two-way ANOVA followed by Šídák’s test for multiple comparisons.

In sum, our data indicate that 5-FU treatment significantly impairs HSC function in a persistent and stem cell-intrinsic manner, resulting in lower engraftment and myeloid-biased output in both PB and BM. Short-term *in vivo* MitoQ treatment alleviates this phenotype, highlighting the translational potential of our findings.

## Discussion

Herein, we have shown using *in vivo* models that repeated exposure to the cytotoxic agent 5-FU induces persistent bone marrow dysfunction. This dysfunction stems from metabolically deflated and transcriptionally deregulated HSCs and cascades down the hematopoietic differentiation hierarchy. Long after exposure, HSCs retain inflammatory signatures, and transplantation assays reveal that these HSCs are stuck in a self-renewing, low-output state, phenocopying the decline of the blood system during natural ageing.

We postulated that the persistent reduction in mitochondrial content and mitochondrial membrane potential of 5-FU exposed HSCs was driving the hematopoietic dysfunction. We thus applied a similar approach, that was previously successful in our hands, to rescue metabolically unfit aged HSCs, by injecting mice with MitoQ. Five days of MitoQ treatment 3 weeks post-chemotherapy was indeed sufficient to significantly correct PB output, HSC transcriptome, and function in a durable manner.

Altered mitochondrial metabolism might not only be a driving force of HSC dysfunction, but may also play a substantial role in CHIP, MDS and AML development (59). Indeed, a recent series of papers highlights the potential for mitochondrial-targeted drugs, including MitoQ and Metformin, to reduce the competitive advantage of premalignant cells (22, 60, 61).

Based on our observations that both natural ageing and cytotoxic treatment establish similar mitochondrial defects in HSCs, we question whether other regenerative stressors, including acute blood loss and viral or bacterial infection, may also manifest their phenotype through lack of mitochondrial recovery in HSCs. If this is the case, similar treatment strategies may be useful.

Although this project addresses fundamental questions in stem cell biology, we are proposing that this work might be broadly therapeutically relevant, if translatable to a human context. First of all, mitochondrial-targeted supportive treatment may accelerate or qualitatively improve hematopoietic recovery between cycles of chemotherapy, improving the quality of life of patients and ensuring treatment continuation. Secondly, we believe that the damaging effect of cytotoxic treatment may have understudied implications for cell therapy, in the context of autologous HSCT with prior conditioning chemotherapy, and in the context of autologous harvest of lymphocytes for cellular immunotherapy, since lymphocytes appear to be most affected by treatment. Lastly, we are intrigued by the possibility that MitoQ treatment might reduce the risk of therapy-related hematopoietic dysplasia (t-MDS) or acute myeloid leukaemia (t-AML) by increasing the overall fitness of the HSC pool.

## Acknowledgements

We thank Jurjen Versluis and Jonas Larsson for their valuable input and discussions at various stages of this project. We thank the animal caretakers of the Erasmus MC animal facility (EDC) for their research support.

This work was funded by the Dutch Cancer Society (KWF: 15215, E.M.), the European Hematology Association (EHA: RG-202012-00206, E.M.) and the Swedish Research Council (VR: 2022-01476, E.M). V.S. was supported by the Swedish Cancer Foundation (CF: 190312PJ, 222348PJ, V.S.) and the Swedish Child Cancer Foundation (BCF: TJ2018-0106, PR2019-0129, V.S.).

## Author contributions

Conceptualization: T.v.T., V.S., E.M.

Investigation and validation: T.v.T., J.L., A.d.S., A.B., W.M., E.B., V.S., E.M.

Methodology: T.v.T., J.L., E.B., V.S., E.M.

Data curation: G.v.B.

Formal analysis: T.v.T., E.M.

Funding acquisition: V.S., E.M.

Software: T.v.T., E.M.

Supervision & Project administration: E.M.

Visualization: T.v.T.

Writing – original draft: T.v.T., E.M.

Writing – review & editing: T.v.T. M.H.G.P.R, T.E., V.S., E.M.

## Declaration of interest

The authors declare no competing interests.

## STAR★ Methods

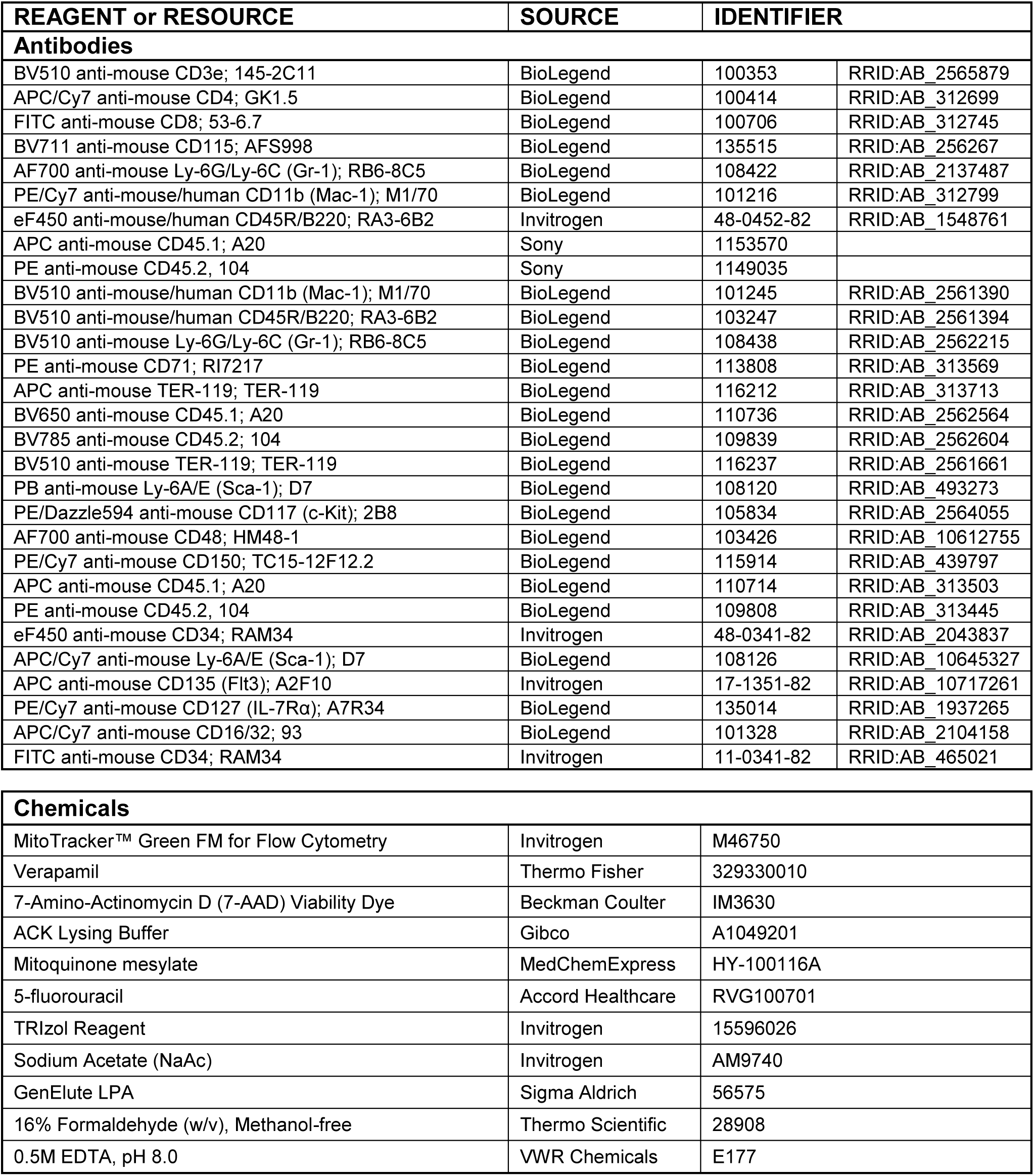
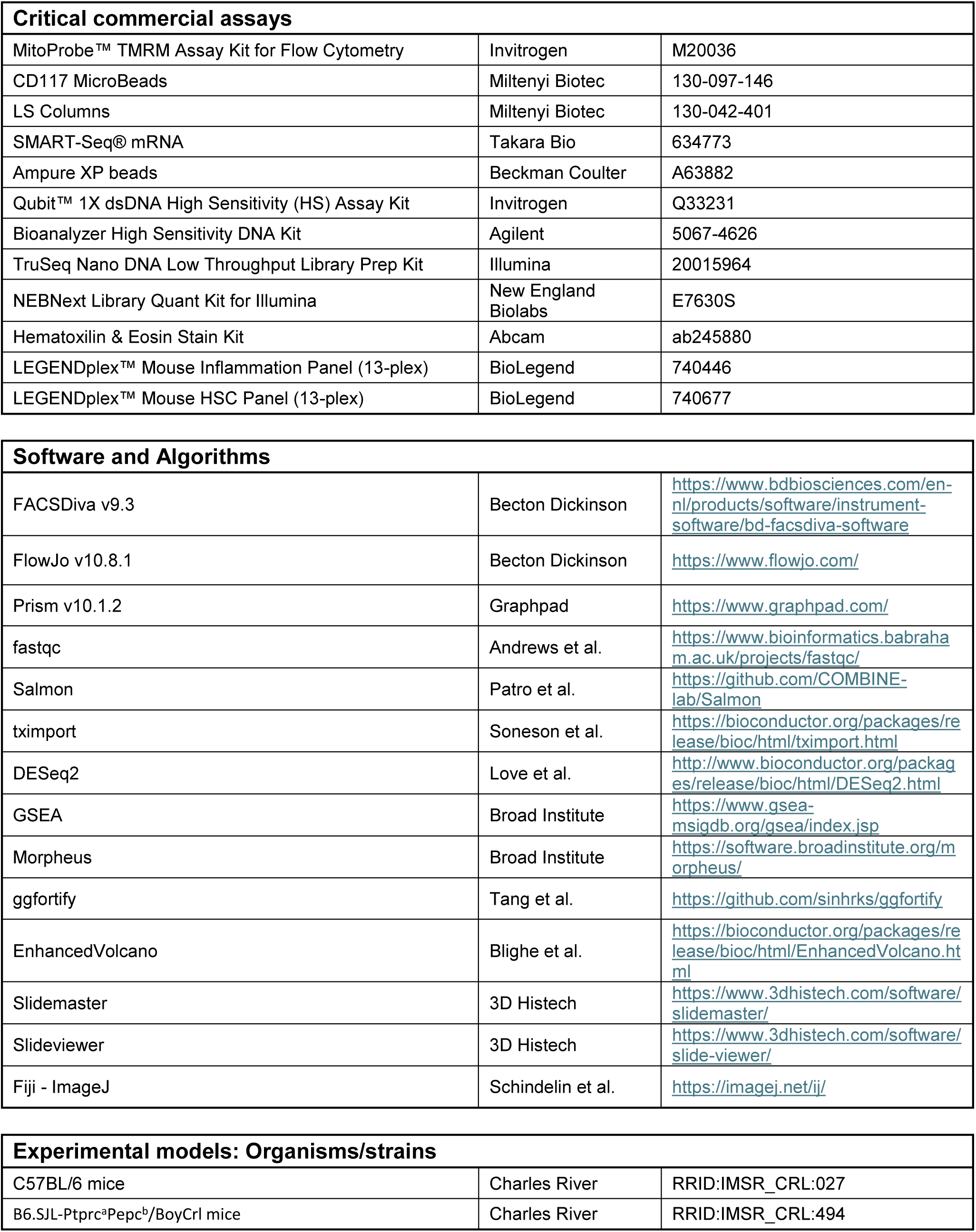

### Mice

Male C57BL/6 mice were purchased from Charles River. All mice were 8-10 weeks old at the time of chemotherapy treatment. Recipients for transplantation were also purchased from Charles River (SJL, B6.SJL-Ptprc^a^Pepc^b^/BoyCrl). All recipient mice were female and 8 weeks old at the time of transplantation. Animals were housed in groups of maximum three animals in individually ventilated cages under specific-free conditions in the Erasmus Animal Center of Erasmus Medical Center in Rotterdam. Chow and water were provided ad libitum and mice were kept in 12-hour light-dark cycles. Experiments and animal care were performed in accordance with the Erasmus Medical Center Animal Ethical Committee.

### Chemotherapy and MitoQ treatment

Male C57BL/6 mice received intravenously (i.v.) injections of 150 mg/kg 5-fluorouracil (Accord Healthcare) diluted in PBS to a final concentration of 25 mg/mL. Mitoquinone mesylate (MedChemExpress, #HY-100116A) was reconstituted in DMSO to 10 mg/mL and diluted in PBS to a final concentration of 0.2 mg/mL before being administered through intra-peritoneal (i.p.) injection at 2 mg/kg body weight.

### Peripheral blood analysis

Peripheral blood (PB) was collected by submandibular bleeding into K_2_EDTA-coated microtainers (Becton Dickinson, #365975). Whole blood count was measured using the scil Vet abc Plus(+) (SCIL Animal Care Company) hematology analyzer. Red blood cells were lysed in ACK lysing buffer (Gibco, #A1049201) at a 1:20 dilution for 5 minutes at room temperature. Lysis was stopped and cells were washed by adding a 10-fold volume of FACS buffer (PBS + 0.5% FCS) prior to staining for cell surface markers.

### Bone marrow analysis

Mice were euthanized by cervical dislocation followed by the dissection of spinal bones and both right and left iliacs, femurs, and tibias. Hind legs and spinal bones were stripped from soft tissue, and bones were crushed in ice-cold FACS buffer using a sterile mortar and pestle. Bone marrow (BM) cell suspensions were filtered (Corning, #431750) and pelleted (500xG, 5 min, 4°C). Red blood cells were lysed in ACK lysing buffer at a 1:20 dilution for 4 minutes on ice. Lysis was stopped and cells were washed by adding a 10-fold volume of FACS buffer prior to staining for cell surface markers. One femur was separated from the remaining bones and processed separately to assess BM cellularity by counting with 1:1 Trypan Blue (Invitrogen, #T10282) using a Countess 3 Cell counter (Invitrogen).

### FACS sorting and analysis

Unless otherwise specified, all antibody incubations were performed at a 1:100 dilution. Except for the erythroid progenitors, all staining procedures were performed on either lysed BM or PB samples. To identify erythroid progenitors, a fraction of whole BM was stained for lineage (CD3e, CD11b, B220, Gr-1), CD71 (1:50) and Ter119 (1:50), on ice for 20 minutes.

To identify differentiated (GMBT) cells, PB and BM cells were stained for CD3e, CD4, CD8, CD115, Gr-1 (1:200), CD11b (1:200), and CD45R (1:200). Cells were stained in FACS buffer for 20 minutes at RT (PB) or 4°C (BM). For hematopoietic stem and progenitor (HSPC) cell analysis, cells were first stained for CD34 (1:50) for 70 minutes at 4°C followed by 20 minutes staining for lineage (CD3e, CD11b, CD45R, Gr-1, Ter119), Sca-1, c-Kit, CD48, CD150, CD16/32, FLT3 (1:40) and CD127 (1:50). Mitochondrial parameters were analyzed by intracellular staining for mitochondrial mass, with MitoTracker Green (MTG, 1:1000, Invitrogen #M46750), and mitochondrial membrane potential, with TMRM (20 nM, Invitrogen, #M20036), in FACS buffer at 37°C for 30 minutes. To prevent efflux of the reagents, cells were simultaneously incubated with verapamil (50 µM, Thermo Fisher #329330010). For hematopoietic stem cell (HSC) sorting, BM was c-Kit enriched using microbeads (1:100, Miltenyi Biotec, #130-097-146) over magnetic columns (Miltenyi Biotec, #130-042-401) before staining for LKS-SLAM cell surface markers (lineage (CD3e, CD11b, CD45R, Gr-1, Ter119), Sca-1, c-Kit, CD48, CD150) at 4°C for 20 minutes.

In all stainings, cells were washed and stained for viability (7AAD, Beckman Coulter, #B88526) prior to analysis or sorting. All centrifugation steps were done at 4°C for 5 minutes at 500xG. When assessing chimerism, markers against CD45.1 and CD45.2 were included. FACS sorting was performed on a BD FACSAria-III, and cells were sorted into TRIzol (Invitrogen, #15596026). When sorting was not required, data was acquired using a BD FACSSymphony A5 and BD FACSDiva Software (BD bioscience, V9.3). Data was processed with FlowJo Software (BD bioscience, V.10.8.1).

### Cytokine quantification in bone marrow supernatant

To obtain bone marrow supernatant, femurs were harvested from individual mice and both ends of the femurs were removed. The bone marrow was flushed from each femur using 300µl PBS and a 30G needle. The resulting cell suspensions were centrifuged at 500xG for 5 minutes at 4°C. The supernatant was carefully collected, snap frozen in liquid nitrogen, and stored at −80°C until further analysis by LEGENDplex^TM^ assays. The LEGENDplex^TM^ assays mouse HSC panel standard (13-plex, Biolegend, #740684) and mouse inflammation panel (13-plex, Biolegend # 740446) were performed following manufacturer’s instructions with minor modifications. In brief, supernatants were incubated overnight at 4°C in an assay buffer:beads:sample ratio of 1:2:1. Beads were measured on low using a BD FACSSymphony A5 and data was analyzed using manufacturers LEGENDplex analysis software (V.2025.05.01).

### Hematoxylin & Eosin staining

Humeri were fixed in 4% paraformaldehyde (Thermo Scientific, #28908) for 24 hours at 4°C, followed by PBS wash and decalcification in 0.5M EDTA (VWR Chemicals, #E177) for 5 days. Decalcified bones were stored in 70% ethanol at 4°C until further processing. Prior to embedding, bones were sequentially dehydrated and paraffinized using a spin tissue processor (Epredia, #813150). Processed bones were embedded in paraffin (Histostar, Fisher Scientific) to generate formalin-fixed paraffin-embedded (FFPE) blocks, which were sectioned at 4 µm thickness. Sections were placed in a 60°C water bath to allow gentle stretching before mounting on standard microscope slides. For staining, slides were deparaffinized in xylene for 5 minutes and rehydrated through a graded ethanol series to a final concentration of 70%. Hematoxylin & Eosin (H&E) staining was performed using the Abcam H&E Stain Kit (Abcam, #ab245880) with minor modifications. In brief, slides were rehydrated in distilled water and incubated in Mayer’s Hematoxylin (Lillie’s Modification) for 8 seconds. Excess stain was removed, with two rinses of distilled water. Nuclear contrast was enhanced by applying bluing agents for 15 seconds, followed by two rinses of distilled water. Prior to eosin staining, slides were briefly immersed (∼2 seconds) in 100% ethanol. Eosin Y solution (Modified Alcoholic) was then applied for 105 seconds. Following staining, slides were rinsed in 100% ethanol and cleared in xylene. Sections were mounted using Entellan (Sigma Aldrich, #1.07961) and glass cover slips. Slides were dried overnight at room temperature before scanning (Pannoramic midi scanner, 3DHistech). Images were independently evaluated for adipocyte content in three separate sessions by a blinded observer.

### RNA sequencing (Low cell number bulk)

Total RNA was extracted from sorted HSCs using TRIzol (Invitrogen, #15596026) and carrier. In brief, chloroform was added to TRIzol at a 1:5 dilution, and total RNA was extracted from the aqueous phase and precipitated by adding equal volume of isopropanol supplemented with 2M NaAc (Invitrogen, #AM9740) and GenElute LPA (Sigma Aldrich, #56575). The RNA pellet was washed twice with 70% ethanol and resuspended in RNAse-free water. cDNA was generated using SMART-Seq v4 Ultra Low input RNA kit for Illumina Sequencing (Takara Bio, #634773) following manufacturer’s instructions. Ampure XP beads (Beckman Coulter, #A63882) were used for size selection and cleanup of PCR products. Quality control of the amplified cDNA was performed using Qubit 3 (Invitrogen, #Q33216) with the High Sensitivity kit (Invitrogen, #Q33231) and BioAnalyzer (Agilent) with the High Sensitivity kit (Agilent, #5067-4626). cDNA was sheared using the Covaris S220 ultrasonicator (Covaris, #500217). 100 ng sheared cDNA was used to generate sequencing libraries using the TruSeq Nano DNA library prep kit (Illumina, #20015964), according to manufacturer’s instructions. Quality control was again performed using both Qubit and Bioanalyzer. Final library concentrations were determined using NEBNext Library Quant Kit for Illumina (New England Biolabs, #E7630S). cDNA libraries were paired end-sequenced (2x 100bp) on the NovaSeq 6000 (Illumina).

### Differential gene expression analysis

Demultiplexing was conducted using the bcl2fastq package (V2.20; Illumina), followed by quality assessment of the resulting FASTQ files using fastqc (v0.11.9) (62). Reads were pseudo-aligned against the mouse reference genome (mm10-build GRCm38.p6) using Salmon (v1.10.0) (63). Transcript quantification files were imported using tximport (v1.36.0) (64) and differential gene expression was performed with DESeq2 (v.1.48.1) in R (V4.5.0) (65). Correction for multiple testing was applied using the Benjamini–Hochberg procedure, and corrected genes with Log2 fold change higher/lower than 0.5 and *P* ≤ 0.05 were considered significantly differentially expressed. Genes were ranked based on adaptive shrinkage (v2.2-54) log2-foldchange, and the resulting ranked lists were stored for downstream analysis (66). Gene-set enrichment analysis (GSEA) was conducted using the Broad-Institute GSEA tool (V4.4.0) (67, 68), employing the classic enrichment score calculation, 1.000 gene-set based permutations, and the signal2noise metric for gene ranking. Analyses were conducted using gene-set libraries from Hallmarks, Reactome, Gene Ontology Biological Processes (GOBP), M5, Mouse Mitocarta (V3.0). Only gene sets containing 15 to 500 genes were included in the analysis. Heatmaps were generated using the web tool Morpheus (broad institute), based on z-scores of log2 transformed transcripts per million (TPM) values *log2(TPM + 1)*. Principle component analysis (PCA) was conducted using the ggfortify (0.4.17) R-package (69), and Volcano plots were created using the EnhancedVolcano (V1.26.0) R-package (70).

### Hematopoietic stem cell transplantation

Recipient SJL mice were lethally irradiated (single dose of 9 Gy) 16-hours prior to transplantation. FACS sorted HSCs from chemotherapy-treated mice (400 HSCs) together with 200.000 whole bone marrow SJL support cells were transplanted in 200µl FACS buffer (PBS + 0.5% FCS) through intravenous tail vein injection. Mice received enrofloxacine-supplemented (0.6 ml/L, Baytril) autoclaved water for 3 subsequent weeks after transplantations to prevent infections during this period of myelo-ablation. Peripheral blood was analyzed every 3-weeks to assess donor engraftment and lineage kinetics. Bone marrow engraftment and lineage kinetics were determined at either 12-week or at the 15-weeks endpoint, as listed in figure legends.

### Quantification and statistical analysis

Statistical analyses were performed in GraphPad Prism 10 (version 10.1.2, Graphpad Software). Results are represented as mean ± S.E.M. For comparisons between only two groups, statistical significance was determined using a two-tailed student’s *t*-test. In case of multiple comparisons to the control group, either a one-way analysis of variance (ANOVA) followed by Dennett’s multiple comparisons test was used or a two-way ANOVA followed by Šídák’s multiple comparisons test, as reported in the figure legends. In case of multiple comparisons between all treatment groups, a two-way ANOVA followed by a Tukey’s multiple comparisons test was used. Asterisks (*P <0.05, **P < 0.01, ***P < 0.001, ****p < 0.0001).

---

**Supplementary figure 1.**

(A) Scil vet ABC plus counter analysis of white blood cells in peripheral blood (PB) of control (N=3, black) and 5-FU exposed (N=6, orange) animals 3 weeks post-chemotherapy treatment.

(B) Percentage granulocyte-monocyte progenitor (GMP, Lineage^-^ C-Kit^+^ Sca-1^-^ CD16/32^+^ CD34^+^) of total BM.

(C) LEGENDplex^TM^ analysis of BM supernatants using the LEGENDplex^TM^ inflammatory panel (upper) or HSC panel (Lower). N=3 mice per group.

(D) Experimental overview.

(E) Left: BM cellularity counts and percentage HSC (Lin^-^ C-kit^+^ Sca-1^+^ CD150^+^ CD48^-^) (Middle) and Long-Term HSC (Lin^-^ C-kit^+^ Sca-1^+^ CD150^+^ CD48^-^ CD34^-^) of total BM in control (N=4, black) and doxorubicin exposed (N=4, red) animals 3 weeks after single injection of Doxorubicin.

(F) Transplantation setup.

(G) Total BM engraftment (Left) and contribution of donor HSC to total HSC pool (Right) in recipients receiving HSCs of control (N=3, black) and doxorubicin (N=3) treated animals 12 weeks post-transplantation.

**Supplementary figure 2.**

(A) Scil vet ABC plus counter analysis of white blood cells (Left), red blood cells (Middle), and platelets (Right) in peripheral blood of control (n=6, black) and 5-FU exposed (N=6, orange) animals, 3, 6, and 12 weeks post-chemotherapy.

(B) BM cellularity counts 12 weeks post-chemotherapy.

Results are represented as mean ± SEM. ^∗^p < 0.05; ^∗∗∗∗^p < 0.0001. Statistical analysis used: unpaired two-tailed student’s *t*-test for two group comparisons and two-way ANOVA followed by Šídák’s test for multiple comparisons.

**Supplementary figure 3.**

(A) Gene set enrichment analysis (GSEA) showing top significantly up- or downregulated pathways comparing HSCs of 5-FU exposed (n=3, orange) and control (n=5, black) animals.

(B) GSEA of mouse.mitocarta (v3) gene set.

(C) Metabolic processes in the Top 20 enriched GOBP GSEA gene set.

**Supplementary figure 4.**

(A) Scil vet ABC plus counter analysis of platelets in peripheral blood (PB) of control + PBS (N=6, black), control + MitoQ (n=4, grey), 5-FU + PBS (N=6, orange) and 5-FU + MitoQ (n=6, green) animals 3, 6, and 12 weeks post-chemotherapy.

Results are represented as mean ± SEM. Statistical analysis used: Ttwo-way ANOVA followed by Dennett’s test for multiple comparisons.

**Supplementary figure 5.**

(A) Volcano plot of differentially expressed genes (DEGs, Log2fold change ≥ 0.5, adjusted *P*-value ≤ 0.05) comparing HSCs of 5-FU + PBS and 5-FU + MitoQ treated animals. Upregulated DEGs are depicted in red and downregulated DEGs in blue.

(B) Heatmap of 28 selected DEGs, which were specifically upregulated or downregulated by MitoQ treatment.

## Notes

### Competing Interest Statement

The authors have declared no competing interest.

