## Supplementary figures and images for "Targeting mitochondria mitigates chemotherapy-induced bone marrow dysfunction"

### Supplementary figure 1

#### Supplementary figure 1

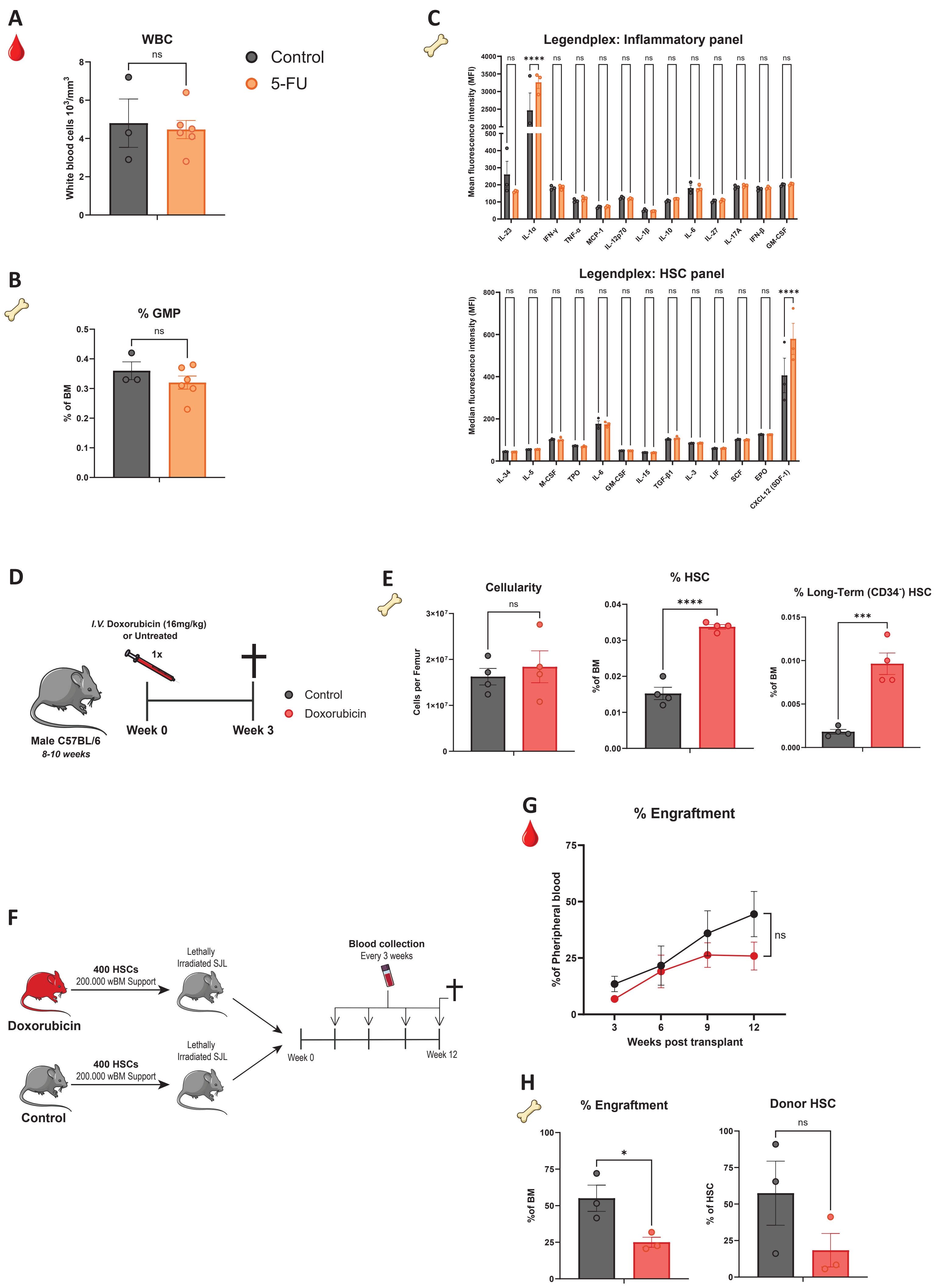

### Supplementary figure 2

#### Supplementary figure 2

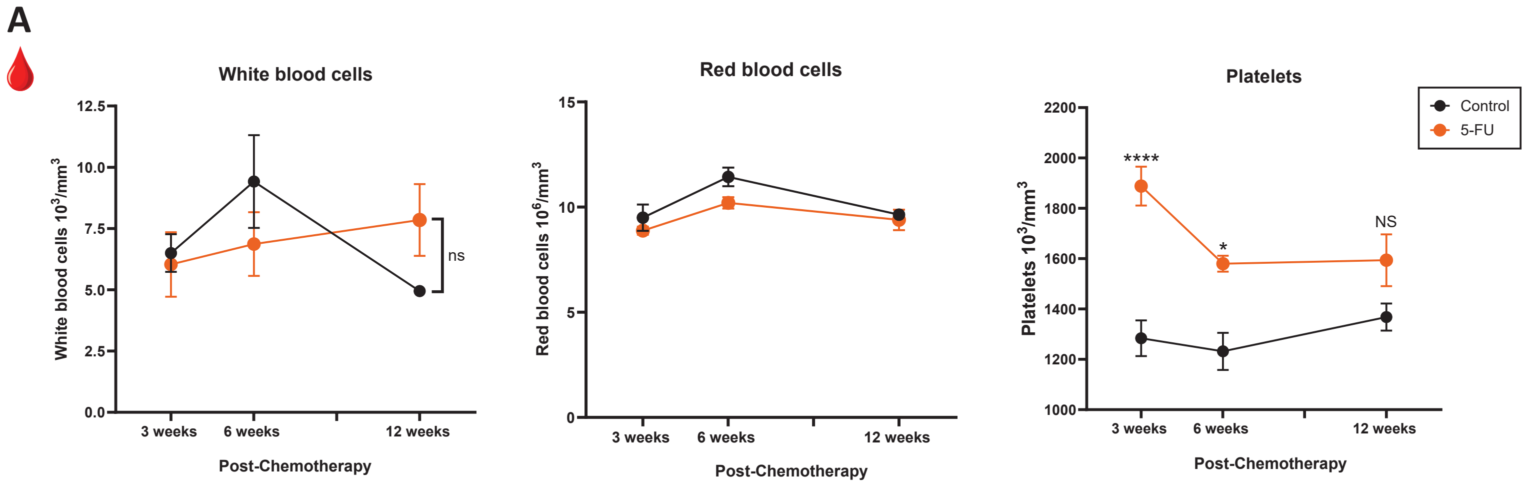


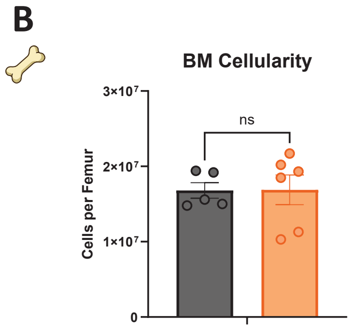

### Supplementary figure 3

#### Supplementary figure 3

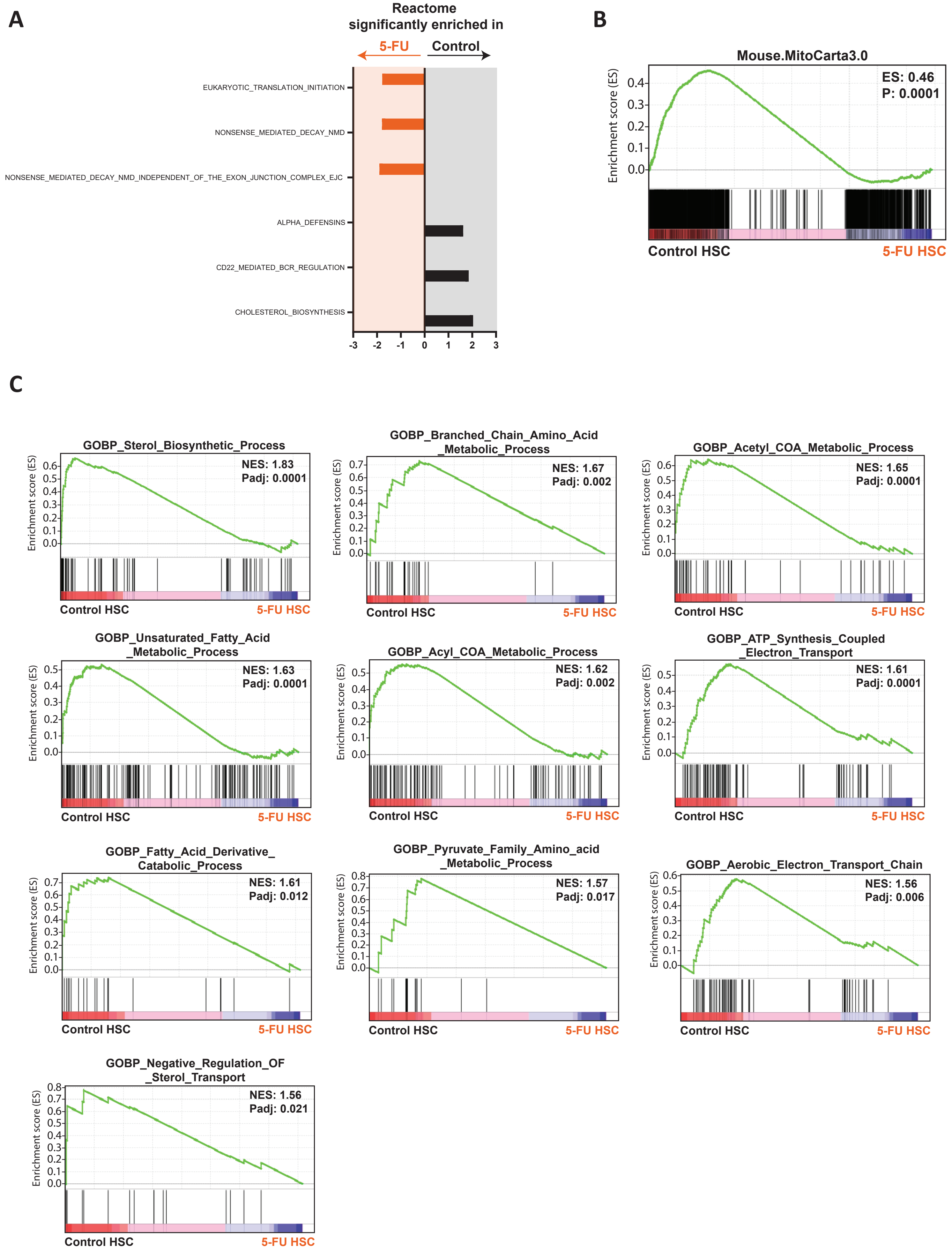

### Supplementary figure 4

#### Supplementary figure 4

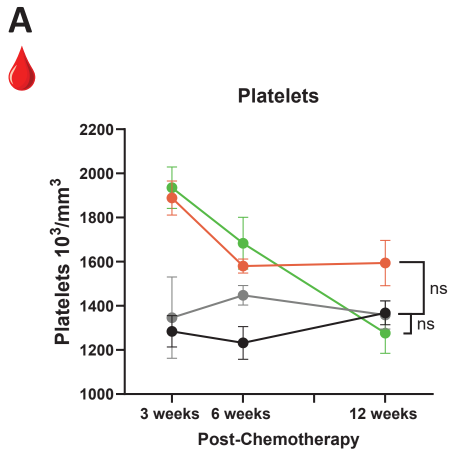

### Supplementary figure 5

#### Supplementary figure 5

A

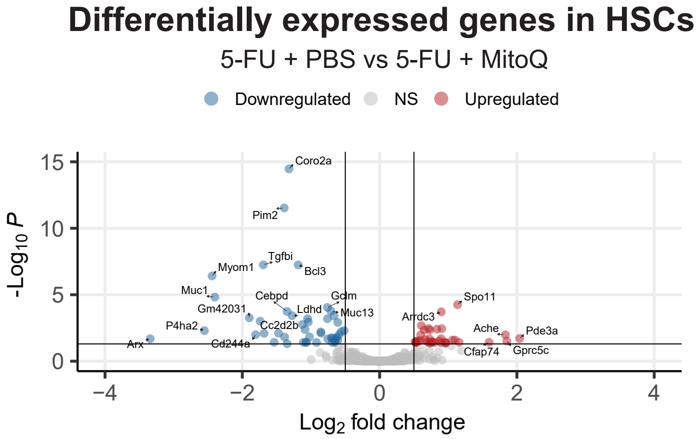


B

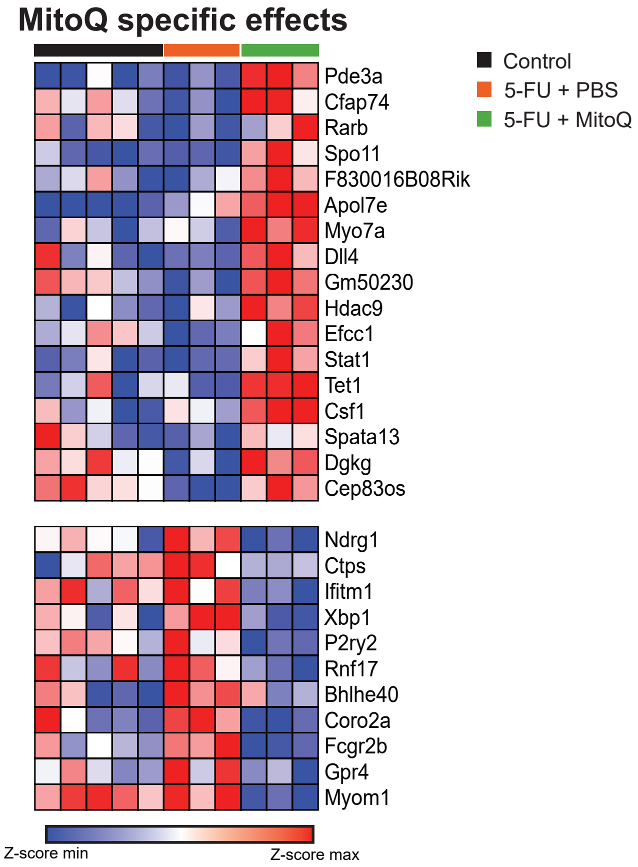
